# Emerging *Mycobacterium bovis* in Lebanon: a snapshot based on whole-genome sequencing

**DOI:** 10.1101/2024.01.18.576209

**Authors:** Israa El Jouaid, Ghena Sobh, George F Araj, Wafaa Achache, Ghiles Grine, Sima Tokajian, Charbel Al Khoury, Fadi Abdel-Sater, Michel Drancourt, Jamal Saad

## Abstract

**Background:** Tuberculosis is a pressing public health issue in Lebanon, a country of approximately five million people, including around 1.5 million refugees from Palestine and Syria. Prior research has revealed uncontrolled animal sources of *Mycobacterium bovis*, emphasizing the necessity for a comprehensive approach to combat tuberculosis in the region.

**Methods:** 48 clinical *Mycobacterium tuberculosis* complex isolates were identified through whole genome sequence. Also, 43 animal fecal samples were collected from various farms across Lebanon to investigate the presence of the *M. tuberculosis* complex using CRISPR-csm4 PCR.

**Results:** Genomic analysis revealed that 39/48 (81.25%) of isolates were *M. tuberculosis* and 9/48 (18.75%) were *M. bovis. M. tuberculosis* was distributed over four lineages, Indo-Oceanic L1 (n = 3/39)(7.6%), East-Asian L2 (n = 1/39)(2.5%), East-African Indian L3 (n = 5/39)(12.8%) and Euro-American L4 (n = 30/39)(76.9%). Sub-lineage L4.8 (Euro-American (mainly T), comprising 8/39 of the isolates (20.5%) was predominant, followed by sub-lineages L3 (East-African Indian, n = 5/39 isolates)(12.8%), L4.2.2.2 (Euro-American (Ural), n= 4/39 isolates)(10.2%) and L4.6.5 (Euro American, n=4/39 isolates)(10.2%). Nine *M. bovis* were classified into two clades, designated as unknown2 (n=2/9; 22.2%) and unknown3 (n=7/9; 77.8%). Interestingly, none of the clades or others were detected in the 48 faecal samples using CRISPR standard PCR and qPCR.

**Conclusions:** This study offers insights into human and bovine tuberculosis in Lebanon, emphasizing *M. tuberculosis* lineages prevalence and *M. bovis* distribution into two clades, aiding the fight against tuberculosis, especially bovine tuberculosis, and renewing our understanding of tuberculosis dynamics in Lebanon.

## Background

Tuberculosis is a significant public health challenge in Lebanon, a Mediterranean country with a population of approximately five million, in addition to an estimated 1.5 million refugees from Palestine and Syria [1–3]. This unique demographic composition and population dynamics have contributed to a notable increase in the incidence and prevalence of tuberculosis within the country. Studies focusing on the *Mycobacterium tuberculosis* [*M. tuberculosis*] complex (MTBC) in Lebanon revealed the presence of abandoned animal reservoirs, particularly for *Mycobacterium bovis* (*M. bovis*) [2, 3]. Limited veterinary control of bovine, caprine and ovine animals, direct exposure to animals including during ritual ceremonies, and the consumption of unpasteurised and undercooked milk, dairy products and other foods, are risk factors for exposure to *M. bovis* in Lebanon [3]. *M. bovis*, with its known for its genetic diversity and comprising 12 genotypes, poses distinctive challenges for tuberculosis management, as 11 of these genotypes are naturally resistant to pyrazinamide, an antitubercular drug commonly used in standard treatment protocols [4]. Despite the recognised importance of *M. bovis* in the epidemiology of tuberculosis, the incidence and prevalence of bovine tuberculosis in Lebanon remain poorly studied. Moreover, there is a growing concern regarding the potential geographic associations between *M. bovis* lineages worldwide, which could have significant implications for public health strategies [4].

To address these knowledge gaps in Lebanon, especially in enhancing understanding of TB and *M. bovis* landscape on the molecular level, this study was warranted. This will be helpful to plan targeted interventions and strategies to combat tuberculosis effectively.

## Methods

### Sample collection

A total of 48 randomly collected clinical isolates, identified as MTBC using BD MGIT™ TBc Identification Test (Becton Dickinson, Sparks, USA) at the Department of Pathology and Laboratory Medicine at the American University of Beirut in Lebanon, were further analysed at the IHU Méditerranée Infection, Marseille, France to confirm the identification at the species level and further characterize the isolates (Table S1). Isolates were anonymised and heat-inactivated prior to shipping to NSB3(Niveau Sécurité Biologique 3) laboratory of IHU Méditerranée Infection. *M. tuberculosis* complex (MTC) isolates were recovered from the following samples: 17 sputum, 12 broncho-alveolar wash (BAL), nine lymph node, two urine, two gastric wash, two tissue, one fine needle aspiration (FNA), one pulmonary fluid, one cerebrospinal fluid (CSF), and abdominal wash (Table S1). Additionally, in December 2022, the French Department for the Protection of Animal and Environmental Health granted IHU Méditerranée Infection prior authorisation to import animal-origin faecal samples into France, allowing for the study of 43 faecal samples from livestock animals, originating from 13 different farms (hosting 269 sheep, 115 cattle, 27 goats and 11 calves) in mount Lebanon (Table 1).

**Table 1.** Faecal samples and collection sites.

| Farms | Type and number of animals | Number of samples | Collected places |
| --- | --- | --- | --- |
| A | 3 cattle, 3 calves, 3 sheep | 5 | Mount Lebanon (Chouf) |
| B | 9 cattle, 4 calves, 7 sheep | 5 | Mount Lebanon (Chouf) |
| C | 20 cattle, 200 sheep, 9 goats | 6 | Mount Lebanon (Chouf) |
| D | 11 cattle, 2 calves, 8 sheep | 6 | Mount Lebanon (Chouf) |
| E | 15 sheep, 18 goats | 5 | Mount Lebanon (Chouf) |
| F | 3 cattle, 2 calves, 6 sheep | 5 | Mount Lebanon (Chouf) |
| G | 30 sheep | 5 | Mount Lebanon (Chouf) |
| H | 10 cattle | 1 | Mount Lebanon (Jbeil) |
| I | 20 cattle | 1 | Mount Lebanon (Jbeil) |
| J | 25 cattle | 1 | Mount Lebanon (Jbeil) |
| K | 5 cattle | 1 | Mount Lebanon (Jbeil) |
| L | 6 cattle | 1 | Mount Lebanon (Jbeil) |
| M | 3 cattle | 1 | Mount Lebanon (Jbeil) |

### DNA extraction

To extract DNA from the collected faecal material, 200 µL of faecal sample mixed with 500 µL of sterile water (UltraPure TM DNase/ RNase-Free distilled water) and glass powder (Sigma Aldrich, St. Louis, MO, USA) was disrupted using a FastPrep apparatus (MP Biomedicals, Santa Ana California, USA) at maximum power for 60 sec. The tubes were then centrifuged at 10 000 g for 1 min and the supernatant, mixed with 180 µL of ATL lysis buffer (Qiagen, Hilden, Germany) and 20 µL of proteinase K (Euromedex, Lyon, France), was incubated for one hour at 56 °C. Following this step, an EZ1 extraction was performed using a Qiagen kit with EZ1 DNA Tissue Kit (Qiagen, Courtaboeuf, France) in accordance with the manufacturer’s guidelines, and the resulting DNA was eluted in a 50 µL volume. A slightly different procedure was followed for clinical isolates which were suspended in 2 mL PBS and subjected to 1 h incubation at 100 °C. 200 µL of the suspension was then mixed with glass powder (acid-washed glass beads 425–600 µm, Sigma-Aldrich, Saint Quentin Fallavier, France), followed by disruption using the FastPrep apparatus at maximum power for 60 sec and centrifugation at 10 000 g for 1 min. The supernatant (300 µL) was mixed with 180 µL of ATL lysis buffer (Qiagen) and 20 µL of proteinase K (Euromedex) and incubated for 1 h at 56 °C. DNA extraction was performed using a Qiagen kit with the EZ1 DNA Tissue Kit following the manufacturer’s instructions (Qiagen) and DNA was eluted in a 50-μL volume [5].

### Standard CRISPR-PCR

PCRs targeting the CRISPR-csm4 gene were performed as previously described [6]. Briefly, PCR amplifications were performed in a C1000™ thermal cycler (Bio-Rad, Hercules, CA, USA) in a 25-μL final volume containing 5 μL of genomic DNA, 12.5 μL AmpliTaq Gold 360 Master Mix (Thermo Fisher Scientific), 0.75 μL of each primer (forward and reverse) and 6 μL of distilled water. The PCR programme included a 15-min denaturation step at 95 °C, then 35 cycles at 95 °C for 30 sec, 60 °C for 30 sec and 72 °C for 90 sec. followed by a final 10 min extension step at 72 °C [6, 7].

### DNA sequencing

Extracted DNA was quantified by a Qubit assay with the high sensitivity kit (Life Technologies, Carlsbad, CA, USA) and 0.2 µg/µL of DNA was sequenced by Illumina MiSeq runs (Illumina Inc., San Diego, USA). DNA was fragmented and amplified by limited PCR (12 cycles), introducing dual-index barcodes and sequencing adapters. After purification on AMPure XP beads (Beckman Coulter Inc., Fullerton, CA, USA), libraries were normalised and pooled for sequencing on the MiSeq. Paired-end sequencing and automated cluster generation with dual indexed 2×250-bp reads were performed during a 40-h run.

### Genome typing and cluster identification

Sequencing reads were analysed using Kaiju with default parameters [8] to determine the contamination level using NCBI BLAST non-redundant protein database, including bacteria, archaea and viruses (2021-02-24 (52 GB)). The overall quality before and after trimming the reads were evaluated using FastQC [9]. Trimmomatic v0.38 [10] was then used to remove residual Illumina adapters and Illumina-specific sequences [10]. Species, lineages and sub-lineages were identified directly using the raw reads through the TB-Profiler (TB_v0.1.3) (https://tbdr.lshtm.ac.uk/) and MTBseq [11] with default settings by mapping to reference *M. tuberculosis* H37Rv (NC_000962.3). MTBseq was used also to recover the statistical mapping data of the reads using *M. tuberculosis* H37Rv (NC_000962.3) as a reference (Table S2). A local SNPs database was used to type *M. bovis* and *M. tuberculosis* Beijing at the sub-lineage level. In addition, a phylogenetic tree was constructed using PhyML v3.3_1 (https://ngphylogeny.fr/tools/), using the *M. bovis* study isolates and 78 reference genomes covering all known clades [4], using the SNPs extracted through MTBseq.

### Detection of drug resistance

The TB-Profiler was used to recover *in silico* susceptibility-resistance profiles to rifampicin, isoniazid, ethambutol, pyrazinamide, streptomycin, fluoroquinolones, aminoglycosides, cycloserine, ethionamide, clofazimine, para-aminosalicylic acid, delamanid, bedaquiline and linezolid. Each detected mutation was confirmed by mapping reads to the reference *M. tuberculosis* H37Rv resistance genes (Table S1).

## Results

### Genome-based Characterization of *M. tuberculosis* Clinical Isolates

Sequence-based analysis revealed that 39/48 (81.25%) isolates identified as *M. tuberculosis* belonged to four different lineages: Indo-Oceanic L1 (n = 3/39) (7.69%), East-Asian L2 (n=1/39) (2.56%), East-African Indian L3 (n= 5/39) (12.82%), and Euro-American L4 (n=30/39) (76.92%) (Fig. 1, Fig. 2, Table S1). The isolates were further divided into 19 sub-lineages. The Indo-Oceanic lineage consisted of sub-lineages L1.1.3.3 (n=1), L1.2.1 (n = 1), L1.2.1.2 (n = 1) and the East-Asian lineage consisted of one sub-lineage L2.2.9 (B0/W148). We detected five isolates with the same East-African-Indian sub-lineage L3, while Euro-American L4 isolates were distributed in 14 sub-lineages: the L4.8 sub-lineage comprised 8/39 of the isolates (20.5%), followed by the L4.2.2.2 (n = 4/39 isolates) (10.25%), the L4.6.5 (n = 4/39) (10.25%), the L4.5 (n=3/39) (7.7%), the L4.3.3 (n=2/39) (5.1%). We also detected one of each of the following sub-lineages: L4.8.1, L4.3.4.1, L4.3.1, L4.2.2.1, L4.2.2, L4.2.1, L4.1.3, L4.1.2.1, L4.1.1.3 (n=1/39) (2.6%) (Fig. 1, Fig. 2, Table S1).

**Fig. 1.**
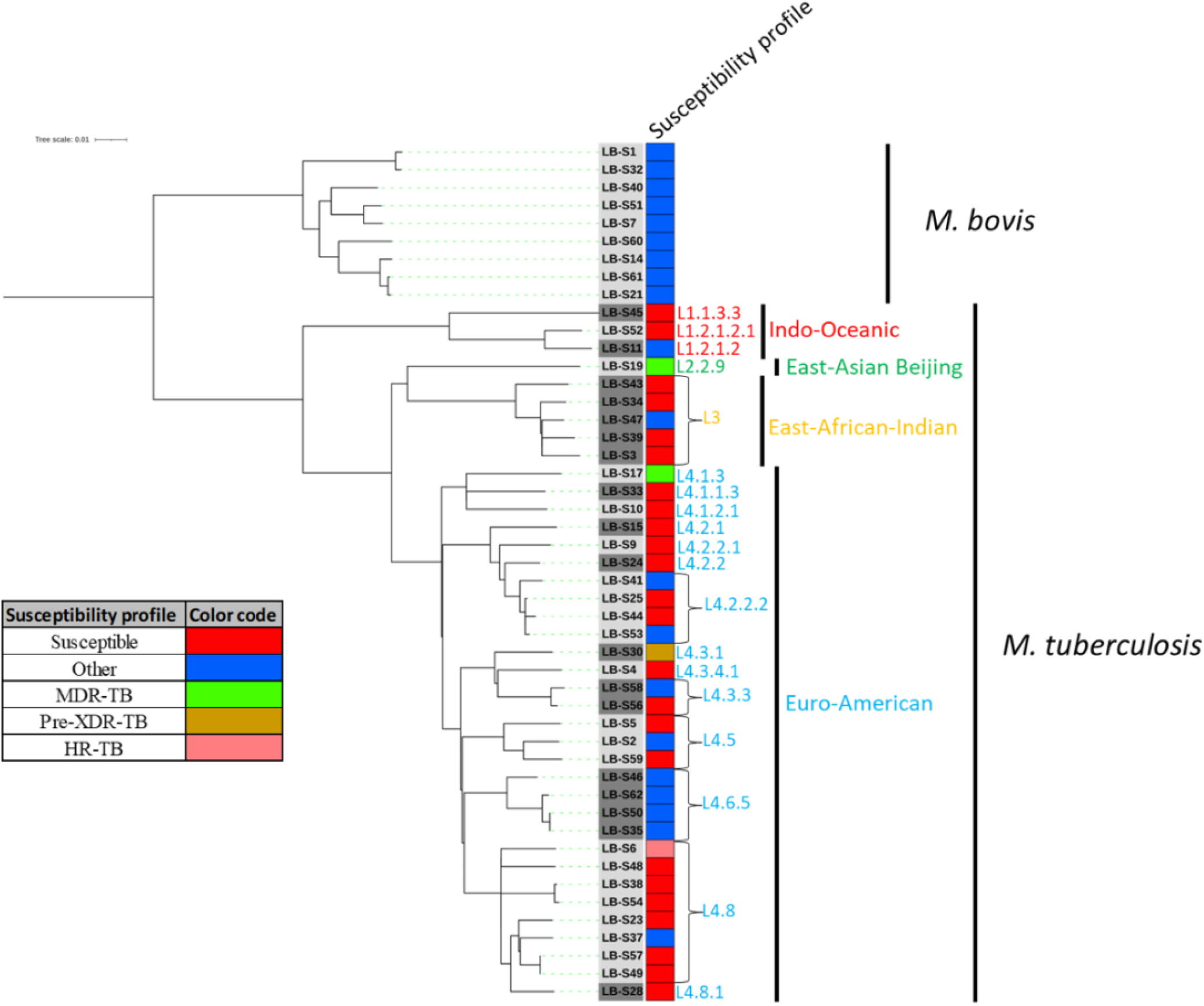
Phylogenetic tree illustrating the diversity within *Mycobacterium tuberculosis* complex in Lebanon. **The** phylogenetic tree was constructed from the aligned set of SNP positions (5918 SNPs) determined by MTBseq in the 48 *Mycobacterium tuberculosis* isolates and nine *Mycobacterium bovis* recovered from Lebanon. The tree was generated using PhyML 3.3.1 software (https://ngphylogeny.fr/tools/) and the GTR model and visualised using the iTOL online website.

**Fig. 2.**
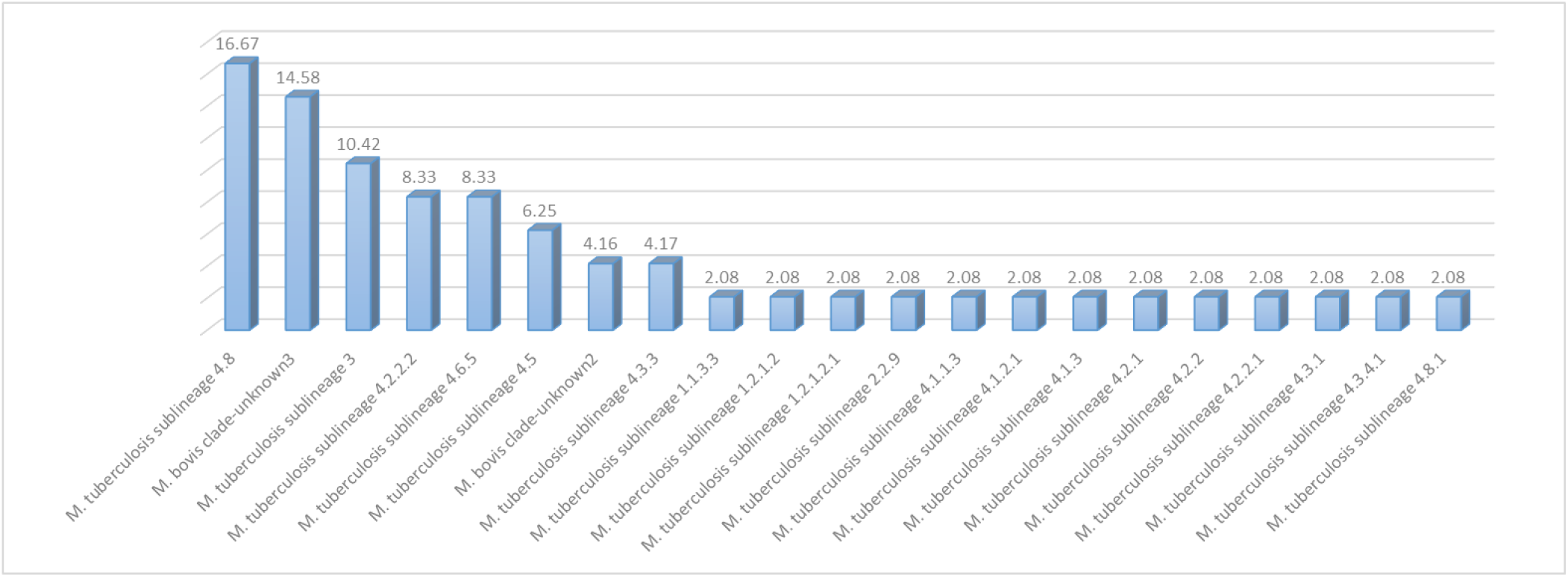
3-D clustered column showing the prevalence in (%) of the *Mycobacterium tuberculosis* sub-lineages (39 isolates) and *Mycobacterium bovis* clades (9 isolates) in Lebanon.

On the other hand, 9/48 (18.75%) studied isolates were identified as *M. bovis* and were divided into two clades: unknown2 (n=2/9) (22.22%) and unknown3 (n=7/9) (77.77%) (Fig. 3, Table S1).

**Fig. 3.**
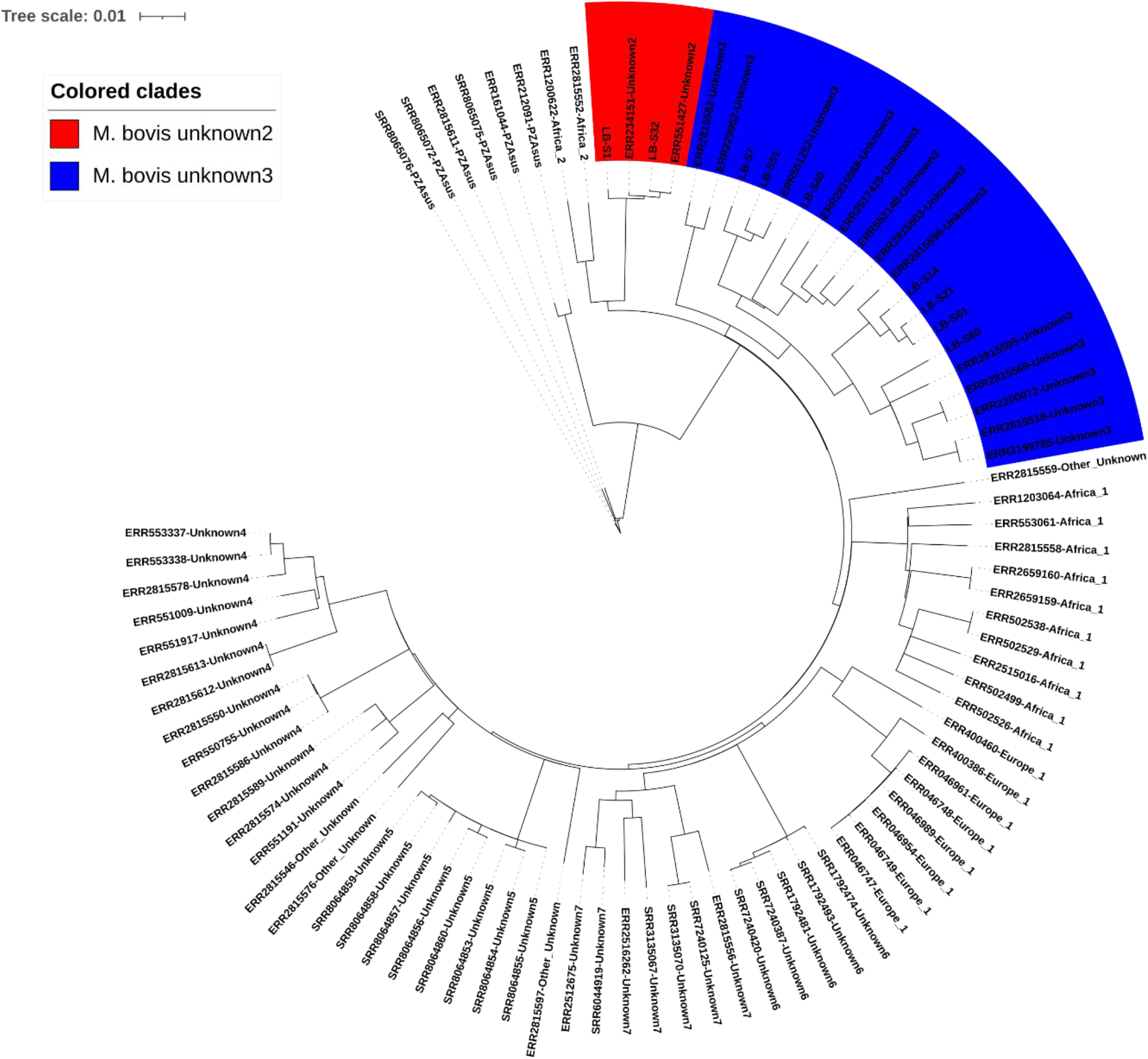
Phylogenetic tree illustrating the clustering of *Mycobacterium bovis* isolates recovered from Lebanon. The Phylogenetic tree was constructed from the aligned set of SNP positions (10 583 SNPs) determined by MTBseq using 87 *Mycobacterium bovis* (nine from Lebanon from this study and the rest from TB-profiler database). Phylogenetic trees were generated using PhyML v3.3.1 software (https://ngphylogeny.fr/tools/) and the GTR model. The tree was visualised using the iTOL online website.

### CRISPR-PCR and qPCR Assays: Environmental MTC Detection

CRISPR-csm4 PCR and qPCR failed to detect any MTC within the 48 faecal samples collected from different animals.

### Antibiotic susceptibility profile

*In silico* antibiotic susceptibility profiling showed that 24/39 (61.5%) of *M. tuberculosis* isolates were susceptible to all tested antibiotics, while 15/39 (38.5%) exhibited at least the anticipated resistance profile (Fig. 1, Table S1). The predicted ones included one pre-extensively drug-resistant (pre-XDR) isolate (LB-S30) belonging to L4.3.1 (n=1/1), two multidrug-resistant (MDR) isolates (LB-S17) belonging to L4.1.3 (n = 1/3) and to L2.2.9 (n=1/1) (LB-S19), and one highly resistant (HR) isolate (LB-S6) belonging to L4.8 (n = 1/7). Eleven *M. tuberculosis* isolates (L1.2.1.2 (n=1/1), L4.3.3 (n=1/2), L4.5 (n=1/2), L4.6.5 (n=4/4), L4.8 (n=1/7), L4.2.2, (n=2/4) and L3 (n=1/5) remained unclassified (termed “Other” by TB-Profiler) (n = 9/15) (60%) and exhibited resistance to one or three antibiotics. All *M. bovis* were only resistant to pyrazinamide (n=9/9) (Fig. 1, Table S1). It is notable that no resistance to linezolid, bedaquiline, delamanid, clofazimine or cycloserine was detected, whereas one *M. tuberculosis* sub-lineage 4.6.5 (LB-S46) showed resistance to para-amino salicylic acid.

## Discussion and conclusion

Drug-resistant tuberculosis is a global health problem causing epidemics worldwide. Whole genome-based characterization in this study provided insights into the drug-resistant tuberculosis in Lebanon. Four human-adapted *M. tuberculosis* lineages were identified: Indo-Oceanic L1, East Asian L2, EastAfrican Indian L3 and Euro-American L4. Notably, the Euro-American L4 lineage was the most prevalent, consistent with previous reports in Lebanon [2, 3]. This lineage also displayed widespread distribution across several North African countries, including Morocco, Tunisia, Algeria, and Libya [12]. Among the other lineages identified, the East African Indian L3 is known to have significant prevalence along the eastern coast of Africa, South India, and Southeast Asia [13]. The Beijing/East Asian lineage L2 has been recognised as a major driver behind the spread of MDR tuberculosis in the Eurasia region [14]. Additionally, the Delhi/Central Asia lineage was observed in the Middle East and Central Asia, with a higher occurrence in India [15].

*M. bovis*, which accounted for 18.75% of the isolates undertaken in this study, was clustered into two distinct clades: “unknown2”, which includes the BCG vaccine strains and was previously found in various regions of East Africa (Eritrea, Ethiopia) as well as Southern Europe (Spain and France) [4], and “unknown3”, a clade distributed across Western Asia and Eastern Europe, with additional presence in East Africa [4]. Looking at it from a broader perspective, data from Lebanese customs authorities (https://al-akhbar.com/In_numbers/304302) for the years 2019 and 2020 revealed that Lebanon imported cattle from various countries, including Spain, Ukraine, Brazil, Hungary, Colombia and Croatia (Table 2). Our genotyping results were in line with the higher prevalence of the “unknown2” and “unknown3” clades in countries such as Spain, Ukraine, and Hungary.

**Table 2.** Number of cattle imported into Lebanon for the years 2019 and 2020. (REF: https://al-akhbar.com/In_numbers/304302).

| Number of cattle imported per year<sup>279</sup> |  |  |
| --- | --- | --- |
| Countries | 2019 | 2020<sup>280</sup> |
| Spain | 50 281 | 49–731 281 |
| Ukraine | 134–986 | 178 700<sup>282</sup> |
| Brazil | 64–167 | 28 594<sup>283</sup> |
| Hungary | 12–270 | 23 271<sup>284</sup> |
| Colombia | 13–521 | 27–387<sup>285</sup> |
| Croatia | 18–570 | 15–707<sup>286</sup> |

Regarding the *M. bovis* resistance profile, this study revealed that all the isolates were resistant to pyrazinamide. It is noteworthy that rifampin, isoniazid, pyrazinamide, and ethambutol are FDA-approved first-line antitubercular medications and are specifically prescribed for the treatment of *M. tuberculosis* infections. Thus, conducting a comprehensive diagnostic assessment is recommended to avoid the use of pyrazinamide for the treatment of *M. bovis* associated infections.

We could not, using the CRISPR approach, detect *M. bovis* in any of the tested faecal material collected from different sites across Lebanon. However, one limitation was the small number of tested samples. Our preliminary results don’t indicate the absence of bovine tuberculosis in Lebanon, especially that previous studies have reported the recovery of *M. bovis* from patients in Lebanon [2, 3]. Accordingly, a larger number of samples collected from more sites especially those with imported animals culd better reflect the status of bovine tuberculosis in the country.

The most commonly used tools in phenotypic and molecular diagnostics are the WHO-approved GeneXpert MTB/RIF test (Cepheid, United States) and matrix-assisted laser desorption ionization-time of flight mass spectrometry (MALDI-TOF-MS) [1]. These diagnostic tools do not differentiate *M. bovis* from the other MTBC species [12]. However, MALDI-TOF-MS method that can be used to differentiate between different strains of the MTBC [16], and which can be used as a diagnostic tool in clinical settings. However, using the MALDI-TOF-MS encompasses the availability of the accompanying databases for the accurate identification of mycobacterial genotypes [16].

Reducing the incidence of tuberculosis in Lebanon requires the implementation of more frequent and better screening strategies to aid in the primary and secondary control and prevention of human and zoonotic tuberculosis. This can be achieved through massive culture-free sequencing of MTBC isolates recovered from animals and clinical samples. Establishing a national database containing metadata and all available DNA sequences will help in better tracking *M. bovis* in dairy cattle products. By monitoring abattoirs, establishing set frequent screening campaigns, and implementing quality control measures will help improve food safety and achieve better public health.

## Ethical declarations

### Ethical approval and consent to participate

Ethical approval was not required as no specific clinical investigation was made for this study which investigated only isolates collected and stored as part of routine clinical care, on a completely anonymised basis.

### Consent for publication

Not applicable.

## Acknowledgements

Not applicable.

## Authors’ contributions

**Conceptualisation:** Fadi Abdel-Sater, Michel Drancourt, Jamal Saad

**Formal analysis:** Israa El Jouaid, Ghena Sobh, George F Araj, Wafaa Achache, Ghiles Grine, Sima Tokajian, Charbel Al Khoury, Fadi Abdel-Sater, Michel Drancourt, Jamal Saad

**Funding acquisition:** Israa El Jouaid

**Investigation:** George F Araj, Ghiles Grine, Sima Tokajian, Charbel Al Khoury, Fadi Abdel-Sater, Michel Drancourt, Jamal Saad

**Project administration:** George F Araj, Ghiles Grine, Sima Tokajian, Fadi Abdel-Sater, Michel Drancourt, Jamal Saad

**Writing – original draft:** Israa El Jouaid

**Writing – review and editing:** George F Araj, Ghiles Grine, Sima Tokajian, Charbel Al Khoury, Fadi Abdel-Sater, Michel Drancourt, Jamal Saad

All authors read and approved the final manuscript.

